# eIF2Bα subcellular localisation – a potential link between translation initiation and stress granule formation?

**DOI:** 10.64898/2025.12.19.695478

**Authors:** Madalena I. Ribeiro de Oliveira, Kelly Parkin-Capper, Susan G. Campbell, K. Elizabeth Allen

## Abstract

The eukaryotic initiation factor 2B (eIF2B) complex is a guanine nucleotide exchange factor (GEF) that regulates translation initiation and is central to the integrated stress response (ISR). eIF2B exists as a decamer composed of catalytic (γ, ε) and regulatory (α, β, δ) subunits, which localise to cytoplasmic foci termed eIF2B bodies. While eIF2B bodies exhibit dynamic changes during stress, the role of individual subunits in the formation of bodies and localisation of subunits to additional cytoplasmic foci remains unclear. Here, we investigate the contribution of eIF2Bα to eIF2B body integrity and investigate alterations in eIF2Bα localisation during cellular stress.

Using immunocytochemistry in neural cell lines, we demonstrate that all endogenous eIF2B subunits form cytoplasmic foci. Whilst eIF2Bα co-localises with the eIF2Bγ catalytic subunit in large bodies (>1 μm²) it also forms smaller foci independently. We find cell type differences in the formation of foci containing eIF2Bα and eIF2Bε subunits, in the absence of changes in protein expression. Stress induction significantly increased eIF2Bα and eIF2Bε foci without altering protein expression, suggesting cell-type specific and ISR-driven reorganisation. siRNA-mediated knockdown of eIF2Bα disrupted large eIF2B body formation, an effect reversed by ISRIB, which stabilizes eIF2B tetramers, highlighting eIF2Bα’s essential role in decamer assembly.

Furthermore, we reveal that eIF2Bα, and to a lesser extent eIF2Bε, localize to SGs under ER and oxidative stress, with colocalisation dependent on eIF2α phosphorylation. Cells harbouring an *EIF2B1* p.Leu99Pro mutation, which impairs ISR sensing, exhibited reduced SG formation and abolished eIF2Bα-SG colocalization. These findings suggest that eIF2Bα not only stabilizes eIF2B bodies but also mediates ISR signalling and SG dynamics via a mechanism reliant on phosphorylated eIF2α.

Collectively, our study uncovers a dual role for eIF2Bα in maintaining eIF2B complex integrity and regulating stress-responsive mRNA triage, providing new insights into translational control and disease mechanisms linked to eIF2B dysfunction.

## Introduction

The effective regulation of protein translation is essential for the maintenance of cellular fitness (Advani & Ivanov, 2019; Hinnebusch et al., 2016). Initiation of translation requires the formation of the ternary complex (TC) between the heterotrimeric eukaryotic initiation factor 2 (eIF2) in its active GTP bound form (eIF2•GTP) and initiator tRNA-Met_i_. The TC together with a host of additional translation initiation factors, binds to the small ribosomal subunit to form the 43S preinitiation complex which then scans the mRNA molecule until it reaches the AUG start codon, upon which translation elongation initiates (Brito Querido et al., 2024; Hinnebusch,2014). Following mRNA start codon recognition, eIF5 GTPase-accelerating protein (GAP) hydrolyses eIF2•GTP resulting in inactive eIF2•GDP on the eIF2γ subunit (Jennings and Pavitt 2010b). The recycling of eIF2 to an active GTP-bound state is essential for continued protein translation, and this process is catalysed by eIF2B via two main functions. eIF2B acts as a GDP dissociation inhibitor, removing eIF5 from the eIF2•GDP/eIF5 complex (Jennings et al., 2013). Additionally, eIF2B acts as a guanine nucleotide exchange factor (GEF) to form the active eIF2•GTP complex (Kenner *et al*., 2019). These eIF2B functions are essential for the formation of ternary complex and continuous rounds of translation.

The integrated stress response (ISR) is a cellular pathway which modulates protein translation in response to stress to maintain cellular homeostasis (Pakos –Zebrucka et al., 2016). Activation of one of four protein kinases phosphorylates eIF2α on serine 51. This phosphorylated eIF2 heterotrimer (eIF2(αP)) binds to and inhibits the GEF activity of the eIF2B protein complex. The lack of active TC due to eIF2B GEF inhibition leads to a global down regulation of protein translation. Concurrently, the translation of a subset of stress-responsive genes is upregulated via the presence of uORFs in the 5’ UTR of the gene (reviewed in Costa-Mattioli & Walter, 2020). In particular, the translational upregulation of the transcriptional activator ATF4 leads to the increased transcription of a range of downstream genes, including GADD34 which leads to the dephosphorylation of P-eIF2α. The ISR aims to maintain or reestablish physiological homeostasis. However, if the stress cannot be resolved, the ISR triggers apoptosis. The ISR is required for memory formation and has been shown to be dysregulated in a range of diseases, from neurodegenerative disease to cancer (Costa-Mattioli & Walter, 2020; Hanson et al., 2022)

The eIF2B protein complex exists as a functional decamer composed of two sets of five non-identical subunits (α-ε), which can be divided into two main categories: catalytic (eIF2Bγ and eIF2Bε) and regulatory (eIF2Bα, eIF2Bβ, eIF2Bδ) subunits (Pavitt et al., 1997). Two eIF2Bβδγε tetramer subcomplexes are stabilised by the eIF2Bα dimer, forming the heterodecamer α2β2γ2δ2ε2 structure (Pavitt et al., 1998; Kashiwagi et al., 2019; Kuhle et al., 2015; Marintchev & Ito, 2020; Schoof et al., 2021; Wortham & Proud, 2015). Additionally, eIF2Bγ and eIF2Bε have been found to form a catalytic subcomplex, which enhances the activity of the ε subunit (Li et al. 2004; Reid et al., 2012).

eIF2B bodies are large assemblies that contain the eIF2B protein complex and were first identified in *S. cerevisiae* and *C. albicans* (Campbell et al., 2005; Egbe et al., 2015; Nuske et al., 2020). In mammalian cells, it was found that different sized eIF2B bodies are present, which vary in their subunit composition. Small bodies appear to be largely comprised of catalytic subunits, whereas medium and large bodies include the regulatory subunits (Hodgson et al., 2019). eIF2 shuttles through eIF2B bodies, and fluorescence recovery after photobleaching (FRAP) analysis has shown a correlation between eIF2B GEF activity and the measurement of eIF2 shuttling (Campbell et al., 2005; Hodgson et al 2019; Norris et al., 2020). In response to stress, the composition of eIF2B bodies changes, suggesting that the presence of eIF2B subcomplexes could have an important role in cellular translation regulation (Wortham et al., 2014; Wortham et al., 2016; Hodgson et al., 2019; Hanson et al., 2024).

The small molecule ISRIB is an inhibitor of the ISR via activation of the eIF2B complex. ISRIB binds a single site on eIF2B, at the interface between the β and δ-subunits of eIF2B (Tsai et al., 2018; Zyryanova et al., 2018). Initial biochemical studies showed that ISRIB stabilises the formation of the full eIF2B decameric complex (Tsai et al., 2018). Crystallisation and cryo EM studies have identified two conformational states of the eIF2•eIF2B complex – the A state and the I state - where A state represents productive eIF2•eIF2B complexes and the I state represents the eIF2B complex bound to one or two eIF2(αP) heterotrimers. Binding of eIF2(αP) creates conformational changes within the eIF2B complex, leading to a change in the binding sites for the eIF2 trimer with the eIF2B complex, steric inhibition of binding of eIF2γ to the catalytic subunits eIF2Bγ & ε, and promotion of binding of a second eIF2(αP) trimer (Schoof et al., 2021; Zyryanova et al., 2021). eIF2(αP) and ISRIB reciprocally oppose each other’s binding to eIF2B, and ISRIB is therefore able to increase eIF2B GEF activity in the presence of eIF2(αP) via antagonistic allostery.

Activation of the ISR impedes the formation of TCs, thereby inhibiting ribosome loading onto mRNA (Jennings et al., 2017). Consequently, polysomes, complexes of actively translating ribosomes along a mRNA, undergo widespread disassembly, releasing a substantial pool of translationally stalled mRNAs into the cytosol which leads to the formation of stress granules (SGs). SGs are phase-separated, membraneless RNP assemblies which act as triage centres for mRNA during periods of cellular stress. SGs execute one of three essential functions: (1) storing translationally silenced mRNA to conserve cellular resources, (2) transporting mRNA transcripts to p-bodies for degradation, or (3) facilitating the transfer of mRNA back into polysomes, thus reinitiating translation when conditions permit (Bley et al., 2015; Campos-Melo et al., 2021; Kedersha et al., 2013). The incorporation of mRNAs into SGs is a tightly regulated process essential for SG assembly and function (Anderson & Kedersha, 2006). Perturbing the equilibrium of mRNA flux by stabilizing polysomes with chemicals like cycloheximide (CHX) or emetine disrupts the formation of SGs induced by stressors that activate the ISR (Hofmann et al., 2021; Glauninger et al., 2022). However, it has also been observed that actively translating mRNA molecules transition between the cytosol and SGs, suggesting that mRNA localization to SGs is compatible with translation (Mateju et al., 2020).

Our studies have identified the formation of eIF2B subcomplexes in cells expressing exogenous eIF2Bε-GFP (Hodgson et al, 2019), however the evidence for the existence of non-decameric eIF2B subcomplexes within the cellular environment has been limited (Zyryanova et al., 2021). We have previously examined the role of eIF2Bα in the formation of eIF2B bodies, and the impact of missense mutations in eIF2Bα on formation of eIF2B bodies in yeast (Norris et al 2020). In this study, we have extended these studies to mammalian cells, to determine the role of eIF2Bα in the formation of eIF2B bodies, and to determine the impact of stress and ISRIB treatment on body formation in the absence of eIF2Bα expression. Our studies aim to investigate further the existence of non-decameric eIF2B subcomplexes in the mammalian cellular environment. Interestingly, we have found an additional localisation property of eIF2B subunits to SGs which give an insight into an additional role of eIF2B subcomplexes during stress.

## Results

### Endogenous eIF2B subunits form cytoplasmic foci in neural cells

Previous analysis of eIF2B body formation and subunit composition utilised exogenously expressed eIF2Bε tagged with mGFP (Hodgson et al 2019; Hanson et al 2024). To analyse endogenous eIF2B localisation, immunocytochemistry of each eIF2B subunit was carried out in three mammalian cell types – U373-MG (astrocytoma), MO3.13 (hybrid primary oligodendrocytes) and SH-SY5Y (neuroblastoma). Neural cell analysis is of particular interest due to differential cell-type stress response and cell-specific contribution to the pathology of vanishing white matter disease, directly linked to mutations in eIF2B subunits (Dooves et al., 2016; Leferink et al., 2019; Hanson et al., 2023; Triñanes-Ramos, et al., 2025). Cytoplasmic foci for all endogenous eIF2B subunits were observed in all cell lines (Fig 1A). Similar proportions of cells formed eIF2B bodies, with 24-29% of cells having localised foci, and the majority of cells displaying a dispersed signal (data not shown).

**Figure 1.**
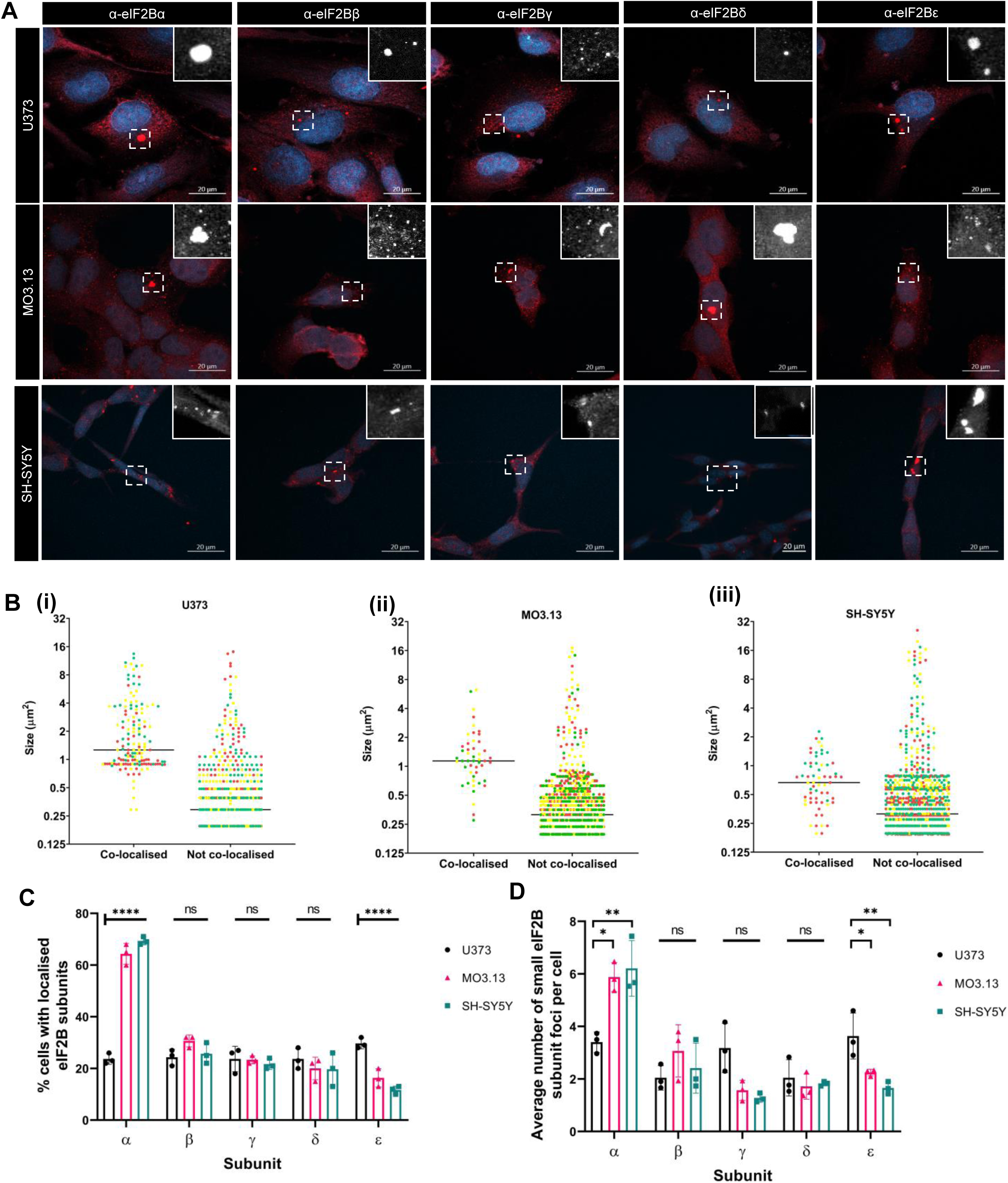
Endogenous eIF2B subunits localise to cytoplasmic foci in a range of neural cell lines. (A) Confocal images of endogenous eIF2B subunits localizing to cytoplasmic foci. U373 and SH-SY5Y cells were fixed in methanol, MO3.13 cells were fixed in 4%PFA and subjected to ICC with (left to right) anti-eIF2Bα, anti-eIF2Bβ, anti-eIF2Bγ, anti-eIF2Bδ, or anti-eIF2Bε primary antibodies and visualized using appropriate secondary antibodies conjugated to Alexa Fluor 594. (B) Co-localised eIF2Bα and eIF2Bγ in U373, MO3.13 and SH-SY5Y (i) – (iii). Size distribution of eIF2Bα foci co-localised and not co-localised with eIF2Bγ (n=3 counts of 30 cells with eIF2Bα localisation). (i) = U373 cells; (ii) = MO3.13 cells; (iii) = SH SY5Y cells (C) Percentage of cells that showed localisation, i.e., one or more foci of eIF2B subunits (n=3 counts of 100 cells) presented as mean ± SD. p Values derived from a two-way ANOVA test, followed by a Tukey’s multiple analysis, *p ≤ 0.05, ***p ≤ 0.001, ****p ≤ 0.0001. (D) Average number of small (<1µm^2^) eIF2Bα, eIF2Bβ, eIF2Bγ, eIF2Bδ, or eIF2Bε foci per cell in U373, MO3.13 and SH-SY5Y cells, presented as mean ± SD (n=3 counts of 30 cells with localised foci). p Values derived from an ordinary one-way ANOVA test, followed by a Tukey’s multiple analysis, *p ≤ 0.05, **p ≤ 0.01. (E) Western Blot analysis of the levels of eIF2Bα expression in U373, MO3.13 and SH-SY5Y cells under untreated conditions. Levels of eIF2Bα were normalized to levels of β-actin. The average ratio of eIF2Bα protein to β-actin is shown under each cell line, standard deviation in brackets

We have shown previously that there is a relationship between eIF2B body composition and size in mammalian cells, with small bodies consisting mainly of catalytic eIF2Bγ and eIF2Bε subunits, and large bodies containing all five catalytic and regulatory eIF2B subunits (Hodgson et al., 2019). For recent studies, we have defined small bodies as <1µm^2^ and large bodies being >1µm^2^ (Hanson et al., 2024) (Fig S1A). When we analysed the size of endogenous eIF2Bα foci, we saw a range of size from 0.2 μm^2^ to 25.2 μm^2^. We reasoned that co-localisation analysis between a catalytic subunit (eIF2Bγ) and a regulatory subunit (eIF2Bα) would allow to distinguish between the localisation of the eIF2B decamer and other subcomplexes. In accordance with previous data, eIF2Bα foci co-localising with eIF2Bγ were of a larger size than non-colocalised eIF2Bα foci (Fig. 1B and S1B).

The proportion of cells displaying subcellular foci of each eIF2B subunit was determined for each cell type. While the formation of eIF2Bβ, γ and δ localised foci did not significantly change between glial and neuronal cells, the proportion of cells which formed localised foci of eIF2Bα and ε varied. SH-SY5Y and MO3.13 cells exhibited a significant increase of percentage of cells with eIF2Bα localisation, whilst U373-MG cells displayed a significantly higher percentage of cells with eIF2Bε foci (Fig. 1C).

When the number of foci of each subunit was determined per cell no significant differences were seen in the number of large foci per cell, whereas significant differences in the number of small foci formed by each eIF2B subunit were observed (Fig 1D and Fig S1C). eIF2Bα formed fewer small cytoplasmic foci in U373-MG cells compared to SH SY5Y and MO3.13 cells, with the reverse being seen with the eIF2Bγ and eIF2Bε subunits (Fig. 1D). This change in the proportion of foci formation was not related to the level of eIF2B subunit expression as determined by western blot (Fig. S1D).

As eIF2B is an essential factor within the cellular ISR, and cellular stress has been shown to change the subunit composition of eIF2B bodies (Hodgson et al., 2019, Hanson et al 2024) we decided to investigate whether the localisation pattern of endogenous eIF2Bα and eIF2Bε changed under ER stress.

For eIF2Bα there was a variable but statistically significant increase in both large and small foci following Tg treatment in all three cell lines (Fig. 2Ai and ii). For eIF2Bε there was no change in the number of large foci, but a significant increase in the number of small foci in all three cell lines following Tg treatment (Fig 2Bi and ii). Western blot analysis showed that the change in number of eIF2Bα and eIF2Bε cytoplasmic foci was not associated with a change in overall protein expression (Fig. 2Ci-iii).

**Fig. 2.**
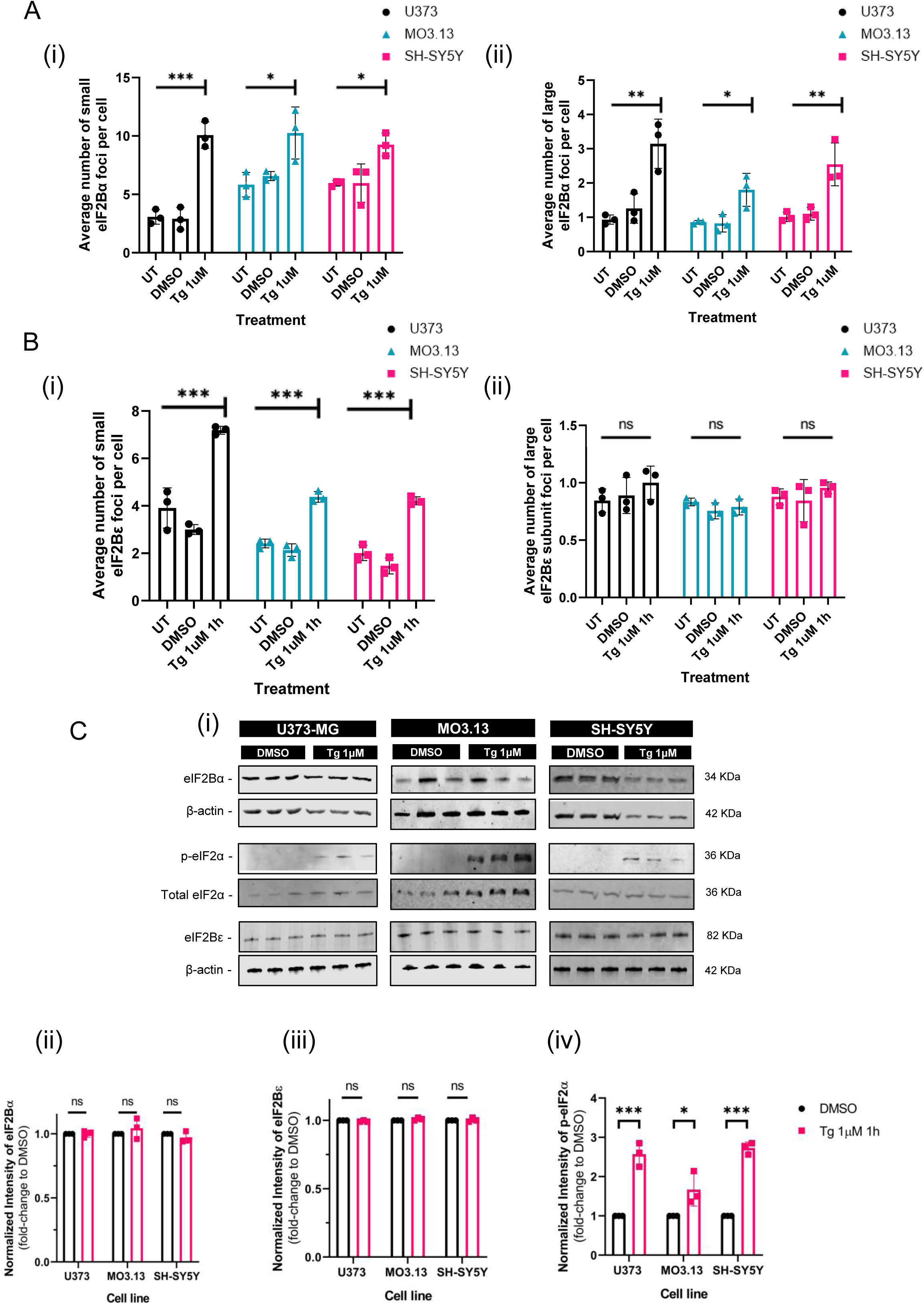
ER stress changes the number of eIF2Bα and eIF2Bε foci. (A) Average number of eIF2Bα foci per cell in a population of 30 cells with localised eIF2Bα, untreated, DMSO and Tg 1μM 1h treatment (N=3). Data was analysed using one-way ANOVA followed by post-hoc Tukey’s test for multiple comparisons. Error bars: ± s.d. (N=3). *p≤0.05; **p≤0.01. (i) small foci <1µm^2^; (ii) = large foci ≥1µm^2^ (B) Average number of eIF2Bε foci per cell in a population of 30 cells with localised eIF2Bε, untreated, DMSO and Tg 1μM 1h treatment (N=3). Data was analysed using one-way ANOVA followed by post-hoc Tukey’s test for multiple comparisons. Error bars: ± s.d. (N=3). *p≤0.05; **p≤0.01. (i) small foci <1µm^2^; (ii) = large foci ≥1µm (C) (i)Western Blot analysis of the levels of eIF2Bα, eIF2Bε p-eIF2α and total eIF2α expression in U373, MO3.13 and SH-SY5Y cells, following DMSO and Tg 1μM 1h treatment. Levels of (ii) eIF2Bα and (iii) eIF2Bε were normalized to levels of β-actin (N=3). (iv) Levels of p-eIF2α were normalized to levels of total eIF2α (N=3). Data was analysed using two-way ANOVA followed by *post-hoc* Tukey’s test for multiple comparisons. Error bars: ± s.d. (N=3). \**p*≤0.05;; \*\*\**p*≤0.001.

Our results show that we can detect the subcellular localisation of all endogenous eIF2B subunits into punctate cytoplasmic foci. eIF2Bα co-localises with the catalytic eIF2Bγ subunit in large cytoplasmic foci (>1µm^2^) and forms smaller foci in the absence of eIF2Bγ. The proportion of eIF2Bα and eIF2Bε foci formed differs between neural cell types, without concomitant changes in the protein expression levels. The number of foci of eIF2Bα and eIF2Bε increases upon ER stress with no significant change in levels of protein expression.

### Knockdown of eIF2Bα protein expression impacts eIF2B body formation, which is reversed upon ISRIB treatment

Missense mutations in the eIF2Bα subunit have been found to impede both decamer formation and the subsequent assembly of eIF2B bodies in yeast (Norris et al., 2021; Taylor et al., 2010). We therefore investigated the impact of the knockdown of eIF2Bα expression on the subcellular localisation of eIF2B subunits and formation of eIF2B bodies.

We found that siRNA knockdown of EIF2B1 reduced eIF2Bα protein expression by 80-95% with no impact on the level of expression of other eIF2B subunits (Fig. 3A & S2).

**Figure 3.**
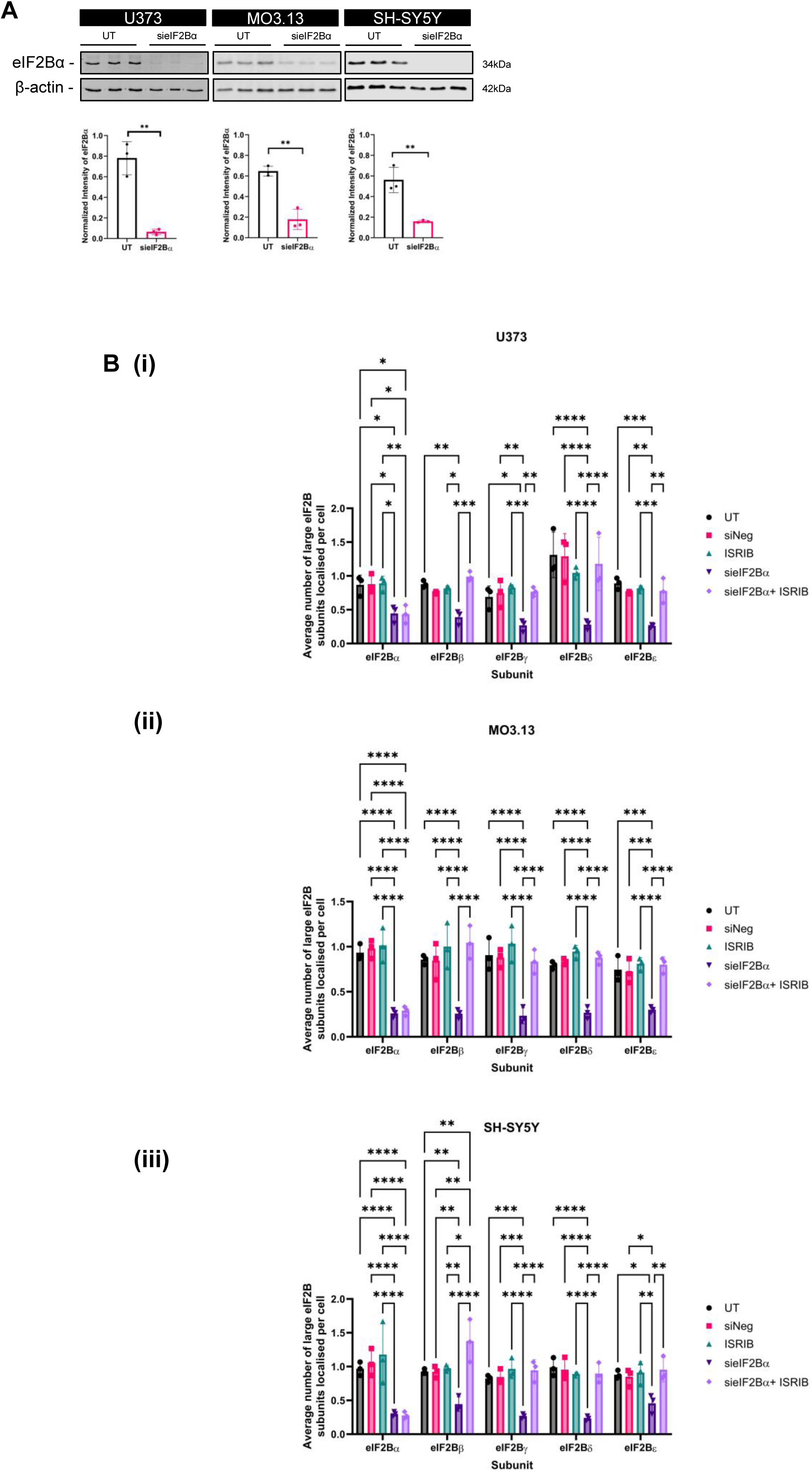
Knockdown of eIF2Bα expression impacts the formation of large eIF2B bodies (≥ 1 μm^2^). (A) Western Blot analysis of the level of eIF2Bα expression in U373, MO3.13 and SH-SY5Y cells following siRNA mediated silencing of eIF2Bα. Levels of eIF2Bα were normalized to levels of β-actin and presented as mean ± SD (n=3). *p* Values derived from an unpaired t test, \*\**p* ≤ 0.01. (B) Average number of large eIF2Bα, eIF2Bβ, eIF2Bγ, eIF2Bδ, or eIF2Bε localized foci per cell following siRNA mediated silencing of eIF2Bα and/or 200 nM ISRIB treatment for 1h, presented as mean ± SD (n=3 counts of 30 cells) in (i) U373 cells, (ii) MO3.13 cells, and (iii) SH-SY5Y cells. *p* Values derived from a two-way ANOVA test, followed by a Tukey’s multiple comparisons analysis, \**p* ≤ 0.05, \*\**p* ≤ 0.01, \*\*\**p* ≤ 0.001, \*\*\*\**p* ≤ 0.0001.

To investigate the impact of lowered eIF2Bα expression on eIF2B body formation, following siRNA mediated silencing of EIF2B1, U373-MG, MO3.13 and SH-SY5Y cells were subjected to ICC targeting the endogenous eIF2Bβ-ε subunits. The knockdown of eIF2Bα did not alter the percentage of cells with eIF2Bβ-ε localised foci in glial and neuronal cell lines analysed (data not shown).

We then went on to investigate the impact of reduced eIF2Bα expression on the number and size of eIF2B bodies. Whilst the impact of reduced eIF2Bα had a slight effect on small eIF2B body formation (data not shown), there was a statistically significant effect on the ability of other eIF2B subunits to be localised to large bodies. We observed that the formation of large eIF2Bβ-ε foci was reduced by eIF2Bα knockdown in glial and neuronal cells. This reduction in the number of large eIF2B bodies formed was reversed when reduction of eIF2Bα expression was coupled with ISRIB treatment (Fig. 3B-iii).

We next explored the impact of the knockdown of eIF2Bα protein expression on ISR activation. Reduction in the expression of eIF2Bα was sufficient to induce the ISR, measured by puromycin incorporation and ATF4 protein expression in the absence of eIF2α phosphorylation, and was rescued by subsequent treatment with ISRIB (Fig. 4A & B).

**Figure 4.**
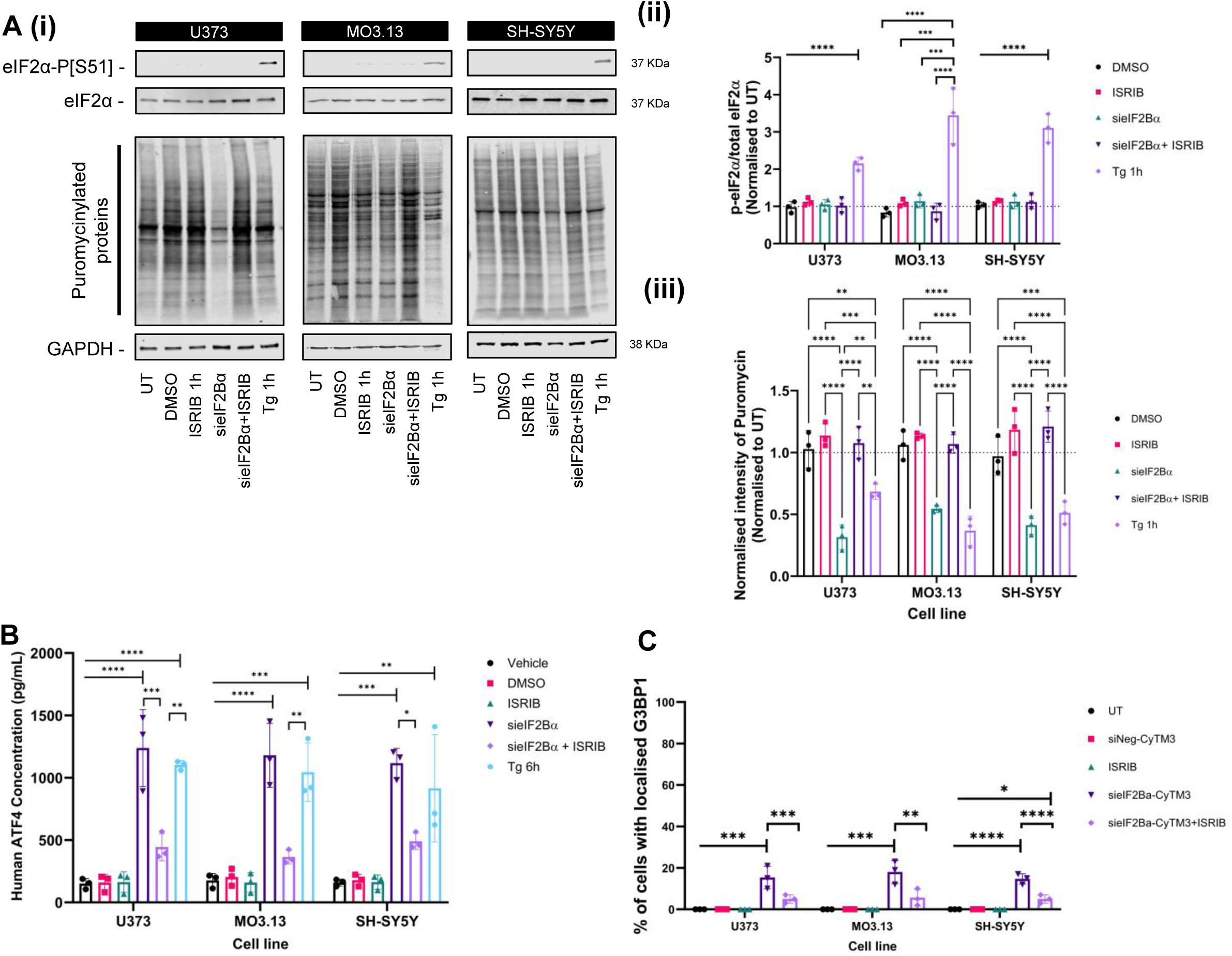
Knockdown of eIF2Bα expression induces the ISR and formation of SG in the absence of P-eIF2α. (A) (i)Western Blot analysis of the level of eIF2α and eIF2α p[S51] expression and puromycin incorporation assays in U373, MO3.13 and SH-SY5Y cells following siRNA mediated silencing of eIF2Bα, 200 nM ISRIB treatment for 1h, or 1 µM Tg for 1h. (ii) Levels of phosphorylated eIF2α were normalized to levels of total eIF2α and presented as mean ± SD (n=3). (iii) Levels of puromycin were normalized to β-actin and are presented as mean ± SD (*n* = 3). *p* Values derived from a two-way ANOVA test, followed by a Tukey’s multiple analysis, \**p* ≤ 0.05, ** *p* ≤ 0.01, \*\*\**p* ≤ 0.001, \*\*\*\**p* ≤ 0.0001. (B) ELISA analysis of the level of ATF4 expression in in U373, MO3.13 and SH-SY5Y cells following siRNA mediated silencing of eIF2Bα, 200 nM ISRIB treatment for 1h, or 300 nM Tg for 6h. Levels of ATF4 detected by ELISA are presented as mean ± SD (n=3). *p* Values derived from a one-way ANOVA test, followed by a Tukey’s multiple analysis, \**p* ≤ 0.05, \*\**p* ≤ 0.01 \*\*\**p* ≤ 0.001, \*\*\*\**p* ≤ 0.0001. (C) Cells were transfected with Cy3 labelled siRNA negative control, Cy3 labelled siRNA targeting EIF2B1, and Cy3 labelled siRNA targeting EIF2B1 coupled with ISRIB 1h (200 nM) treatment. U373-MG and SH-SY5Y cells were fixed in methanol, MO3.13 cells were fixed in 4%PFA and subjected to ICC with anti-G3BP primary antibody and visualized using appropriate secondary antibodies conjugated to Alexa Fluor 488. Mean percentages of U373-MG, MO3.13 and SH-SY5Y cells with G3BPcontaining SGs. Error bars: ±s.d. Data was analysed using one-way ANOVA followed by a Tukey’s multiple analysis. *p ≤ 0.05, **p ≤ 0.01, ***p ≤ 0.001, ****p ≤ 0.0001

The knockdown of expression of eIF2Bα also induced the formation of SGs in 15-20% of cells. This result is surprising given that a previous functional RNAi screen identified that EIF2B1 expression is required for the assembly of SGs, whereas no other eIF2B subunits were identified in this functional screen (Ohn et al., 2008). However, we note that the number of cells expressing stress granules upon Tg treatment is much higher, where over 80% of cells express stress granules (compare Fig. 4C with Fig. 7B).

We hypothesise that the loss of eIF2Bα reduces the proportion of the full eIF2B decameric complex within the cell, and therefore a reduction in formation of large eIF2B bodies. This reduction in formation of the full decameric eIF2B complex induces the ISR, presumably due to a reduction in the GEF activity of the eIF2B complex. Treatment with ISRIB stabilizes what we surmise to be active eIF2B(βγδε)2 structures.

### eIF2B subunits colocalise with stress granules

Whilst these knockdown results show the importance of eIF2Bα in the assembly of eIF2B bodies containing eIF2B regulatory and catalytic subunits, they do not explain the biological function (if any) of the subcellular localisation of eIF2Bα into cytoplasmic foci in the absence of the catalytic eIF2Bγ subunit (Fig. 1B). It has been shown that eIF2Bα can form homodimers (Bogorad et al., 2014; Wortham et al., 2014), and concatemerisation of these dimers may form structures which appear as punctate cytoplasmic foci upon our ICC analysis.

To determine the mechanism causing changes in eIF2B subunit localisation following stress, we determined the colocalisation of eIF2Bα to other membraneless cytoplasmic granules. No colocalisation of eIF2Bα to P bodies was seen (data not shown). Due to a previous study indicating a requirement of *EIF2B1* expression for SG formation (Ohn et al., 2008), we next determined whether eIF2Bα and eIF2Bε subunits localised with SGs formed in response to P-eIF2α. Neuronal and glial cell lines were subjected to ER stress (1 hour Tg treatment at 1 μM), mild oxidative stress (30 minutes sodium arsenite (SA) treatment at 125 μM) and strong oxidative stress (1 hour SA treatment at 500 μM). Cells were subjected to ICC with antibodies to G3BP1, a SG core marker, and eIF2Bα, followed by confocal microscopy analysis. We found that all cell lines showed a significant amount of co-localisation between eIF2Bα and SGs. 24-52% of eIF2Bα and G3BP1 foci co-localised under different stress conditions in U373, MO3.13 and SH-SY5Y cells (Fig. 5A & S3). When the same cells were examined for eIF2Bε and G3BP1 co-localisation after 1 hour SA treatment at 500 μM, 3-21% of eIF2Bε and G3BP foci co-localised (Fig. 5B). Airyscan imaging was performed on eIF2Bε, eIF2Bα and G3BP foci. Partial co-localisation between eIF2Bε and G3BP foci was seen. whereas eIF2Bα and G3BP entirely spatially overlapped (Figure 5C & S3).

**Fig. 5.**
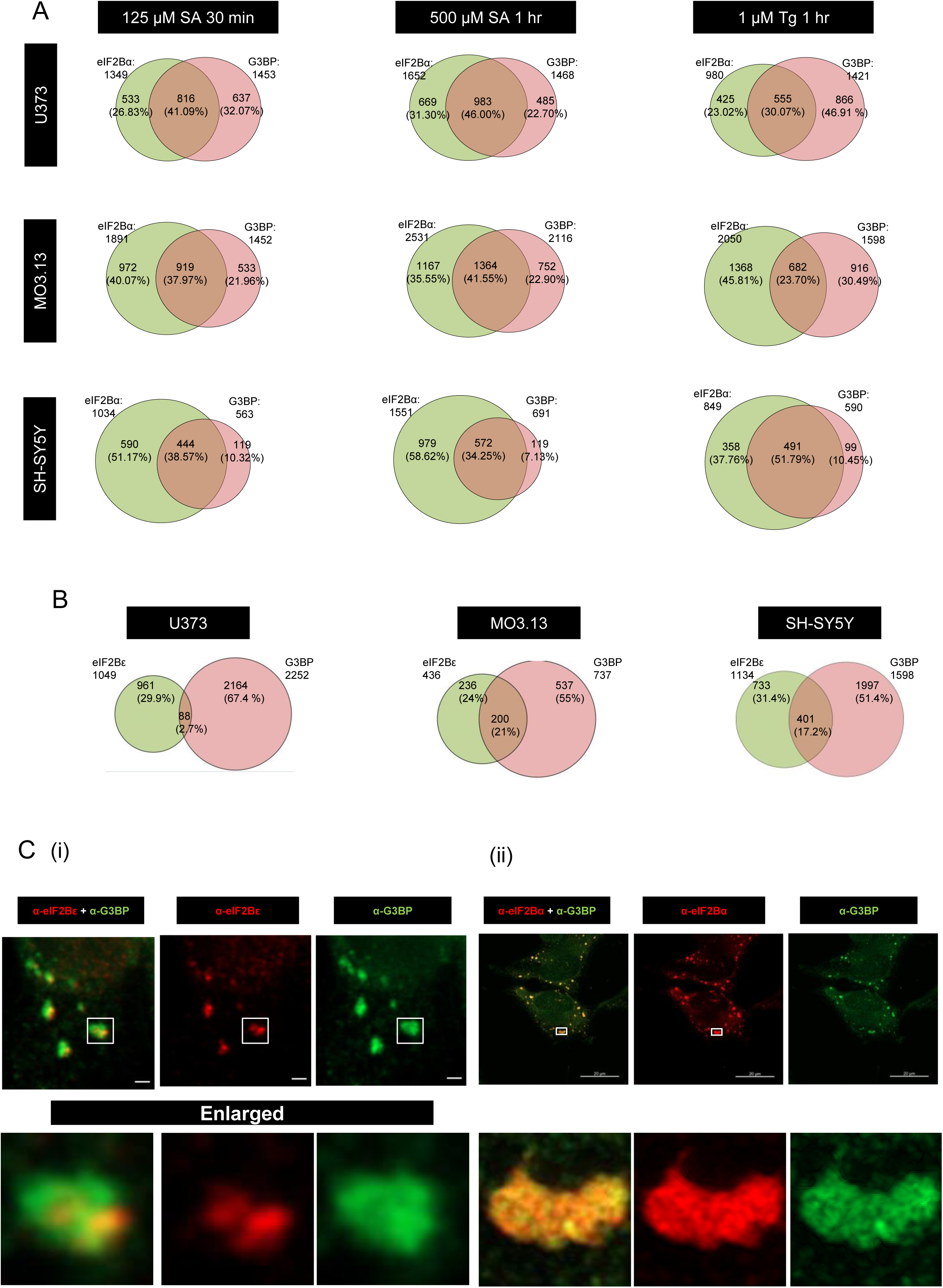
Colocalisation analysis of eIF2Bα and eIF2Bε with ISR induced SGs. (A) Cells were treated with 125 μM SA for 30 minutes, 500 μM SA for 1h or 1 µM Tg for 1h and subjected to ICC with anti-eIF2Bα (green) and anti-G3BP1 (red) primary antibodies visualized using appropriate secondary antibodies conjugated to Alexa Fluor 488 and 594. Venn diagram shows total number of eIF2Bα and G3BP1 foci with the number of foci showing co-localisation (n=3 counts in 30 cells with eIF2Bα localisation). (B) Cells were treated with 500 μM SA for 1h and subjected to ICC with anti-eIF2Bε (green) and anti-G3BP1 (red) primary antibodies visualized using appropriate secondary antibodies conjugated to Alexa Fluor 488 and 594. Venn diagram shows total number of eIF2Bε and G3BP1 foci with the number of foci showing co-localisation (n=3 counts in 30 cells with eIF2Bε localisation). (C) Representative Airyscan images of (i) eIF2Bε and G3BP and (ii) eIF2Bα and G3BP following SA 1h (500 μM) treatments in SH-SY5Y and U373 cells respectively. eIF2Bε panel scale bar: 1 μM. eIF2Bα panel scale bar: 20 μM.

Our results show that eIF2B subunits relocalise to SGs under conditions of ER and oxidative stress. It appears that there is a stronger association between SGs and eIF2Bα than between SG and eIF2Bε.

### Colocalisation of eIF2Bα with SG is dependent on phosphorylation of eIF2α

eIF2Bα is essential for the response to the phosphorylation of eIF2α, making contact with the eIF2α subunit in its phosphorylated form (Schoof et al., 2021; Zyryanova et al., 2021). It has been shown that P-eIF2α is present within SGs (Kedersha et al., 2002). We therefore wanted to investigate whether the co-localisation of eIF2Bα to SGs is dependent on an interaction between eIF2Bα and P-eIF2α. We utilised an ovarian adenocarcinoma SK-OV-3 cell line bearing a heterozygous *EIF2B1* p.Leu99Pro missense mutation, and its wild-type counterpart. The *EIF2B1* p.Leu99Pro mutation is a helix breaker mutation situated within the same helix as several amino acids which make direct contact with P-eIF2α (De Franco et al., 2020) (Fig 6D). This heterozygous *EIF2B1* p.Leu99Pro missense change has been shown to be unresponsive to ISR activation (Powers, Clarke, et al., unpublished data).

**Fig. 6.**
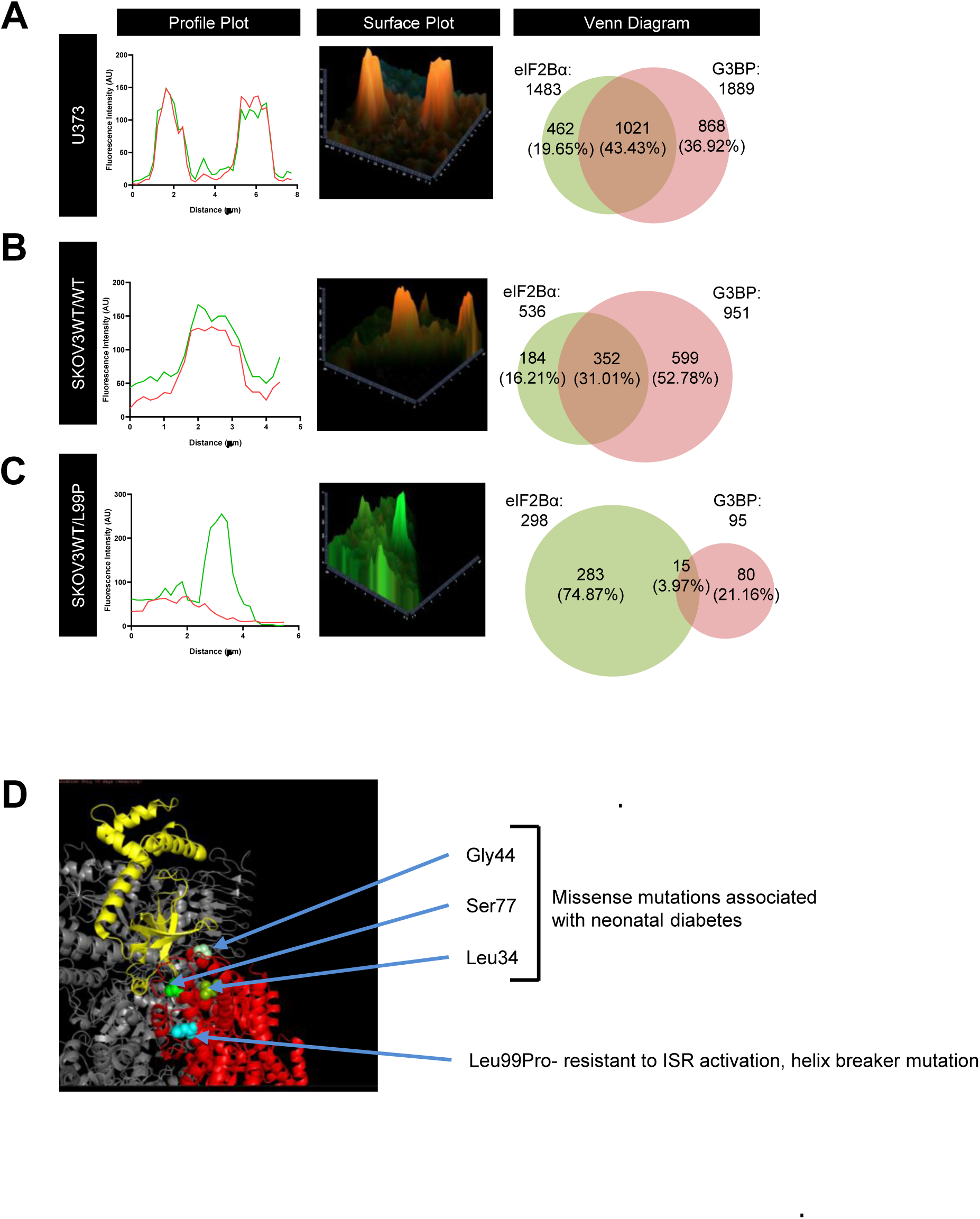
A heterozygous p.Leu100Pro mutation in eIF2Bα disrupts colocalisation of eIF2Bα with ISR induced SGs and impacts the formation of SGs. Analysis of endogenous eIF2Bα and G3BP1 localisation to cytoplasmic foci in (A) U373; (B) SKOV3 *EIF2B1*^WT/WT^ and (C) SKOV3 *EIF2B1*^WT/L100P^ cells following treatment with 500 μM SA for 1h. Cells were subjected to ICC with anti-eIF2Bα (green) and anti-G3BP1 (red) primary antibodies and visualized using appropriate secondary antibodies conjugated to Alexa Fluor 488 and 594. Profile and surface plots of a representative focus show the level of colocalization (separate colours shown on graphics). Percentage of foci are presented as mean ± SD (n=3 counts in 30 cells with eIF2Bα localisation). Venn diagram of eIF2Bα and G3BP populations and co-localisation (n=3 counts in 30 cells exhibiting eIF2Bα localisation). (D) Pymol diagram showing the interaction between eIF2Bα (red chain) and P-eIF2α (yellow chain). Missense mutations associated with neonatal diabetes (Franco et al., 2020) and the mutation present in the SKOV3^WT/L99P^ cells are highlighted

U373, SKOV3 *EIF2B1*^WT/WT^ and SKOV3 *EIF2B1*^WT/L99P^ cell lines were treated with SA (1 hour 500 μM SA treatment), subjected to ICC with antibodies to G3BP1 and eIF2Bα, and analysed by confocal microscopy (Fig. S4). U373 and SKOV3 *EIF2B1*^WT/WT^ cells showed a similar degree of colocalisation of eIF2Bα with SGs (Fig. 6A & B). In contrast, SKOV3 *EIF2B1*^WT/L99P^ cells showed no significant co localisation of eIF2Bα with SGs and also had a reduction in the number of SGs formed upon SA treatment (Fig. 6C, 4 % colocalization; 95 G3BP1 foci formed in 90 cells).

To investigate further the role of P-eIF2α in the colocalization of eIF2Bα with G3BP1, cells were subjected to stress conditions known to cause the formation of SG in a P-eIF2α independent manner. U373, SKOV3 *EIF2B1*^WT/WT^ and SKOV3 *EIF2B1*^WT/L99P^ cell lines were treated with 0.5 mM H_2_O_2_ or 500 nM RocA for 1 hour, subjected to ICC with antibodies to G3BP1 and eIF2Bα, and analysed by confocal microscopy (Fig. S4, Fig7A). In all 3 cell lines, H_2_O_2_ or RocA treatment induced the formation of SGs, however a much lower proportion of SG showed eIF2Bα colocalization (Fig. 7A).

**Fig. 7.**
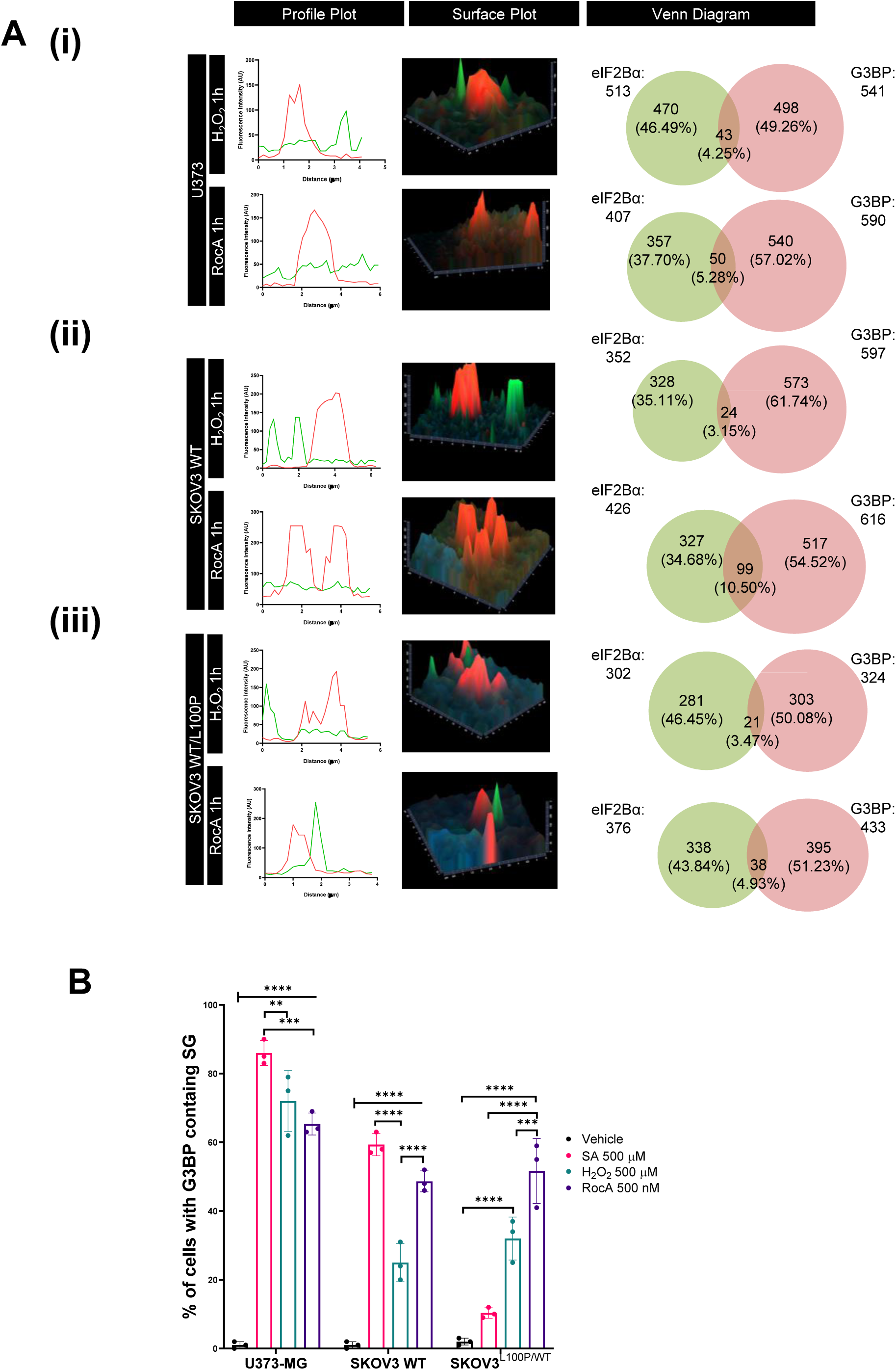
eIF2Bα foci do not colocalise with SG formed via P-eIF2α independent pathways. A. Analysis of endogenous eIF2Bα and G3BP1 localizing to cytoplasmic foci in (i) U373; (ii) SKOV3 *EIF2B1*^WT/WT^ and SKOV3 *EIF2B1*^WT/L100P^ mutant cells following treatment with, 0.5mM H_2_O_2_ or 500 µM RocA for 1h. Cells were fixed and subjected to ICC with anti-eIF2Bα (green) and anti-G3BP1 (red) primary antibodies and visualized using appropriate secondary antibodies conjugated to Alexa Fluor 488 and 594. Profile and surface plots of a representative focus show the level of colocalization (separate colours shown on graphics). Percentage of foci are presented as mean ± SD (n=3 counts in 30 cells with eIF2Bα localisation). Venn diagram of eIF2Bα and G3BP populations and co-localisation (n=3 counts in 30 cells with eIF2Bα localisation). B. Percentage of U373, SKOV3 EIF2B1^WT/WT^ and SKOV3 EIF2B1^WT/L100P^ cells with G3BP containing stress granules following 500 μM SA for 1h, 0.5mM H_2_O_2_ or 500 µM RocA for 1h treatments (n=3 counts of 100 cells) . Mean percentage of cells displaying G3BP-containing SGs in a population of 100 cells per repeat. Error bars: ± s.d. (n=3). Data was analysed using one-way ANOVA followed by a Tukey’s multiple analysis. **p ≤ 0.01, ***p ≤ 0.001, ****p ≤ 0.000

A comparison of the percentage of cells which formed SGs following treatment with SA, H_2_O_2_ and RocA illustrated that there was a significant decrease in the percentage of SKOV3 cells harbouring the *EIF2B1* p.Leu99Pro mutation forming SG when compared to the SKOV3 parent cell line (Fig. 7B)

Collectively, this data shows that the colocalisation of eIF2Bα to SG is impacted either by the inability of eIF2Bα to respond to the phosphorylation of eIF2α, or by alternative stresses which induce the formation of SG in a P-eIF2α independent manner. Additionally, the proportion of cells forming SGs following induction of the ISR by oxidative stress is reduced in cells carrying the p.Leu99Pro mutation in the *EIF2B1* gene, which abolishes the ability of cells to sense the ISR.

## DISCUSSION

### Subcellular localisation of eIF2B subunits may represent formation of key eIF2B subcomplexes

Despite the significant strides made in understanding eIF2B decamer structure (Gordiyenko et al., 2014; Kashiwagi et al., 2016; Schoof et al., 2021; Tsai et al., 2018; Wortham et al., 2014), the precise mechanism of eIF2B complex assembly within the cell remains a subject of ongoing debate. A growing body of evidence suggests that the assembly process of the eIF2B decamer is orchestrated through precursors of sub-complexes (Wortham et al., 2016). This process begins with the initial formation of eIF2Bγε heterodimers, governed by their dimerization capacity. Subsequently, heterodimers of eIF2Bβ and δ form, which then bind to the existing eIF2Bγε heterodimers, culminating in the emergence of an intermediate eIF2Bβδγε tetramer. MS and crystallography results showed that interactions between eIF2Bε-β and γ-δ subunits primarily stabilize the tetrameric subcomplex (Kashiwagi et al., 2016; Wortham et al., 2014). eIF2Bα exhibits the capacity to form homodimers, which in the final stage of eIF2B assembly could bind two tetramers to generate the fully mature eIF2B(αβδγε)_2_ holocomplex (Wortham et al., 2016). Our observations of the localisation of eIF2B subunits may represent the subcellular organisation of eIF2B into cytoplasmic bodies representing stages in this assembly process.

Firstly, we analysed the subcellular localisation of the eIF2Bα and eIF2Bγ subunits to represent the localisation of full eIF2B decamers because eIF2Bα and eIF2Bγ have not been shown to form subcomplexes in the absence of other eIF2B subunits (Kuhle et al., 2015; Pavitt et al., 1998; Schoof et al., 2021). Our initial experiments showed that eIF2Bα co-localises with the catalytic eIF2Bγ subunit in cytoplasmic foci greater than 1µm^2^, but also forms foci in the absence of eIF2Bγ. The subunit composition of the larger foci agrees with our previous data (Hodgson et al., 2019).

Further investigation of the subcellular localisation of eIF2B subunits showed that the punctate localisation of eIF2Bα and eIF2Bε varies between cell types (Fig. 1). Treatment of cells with agents that activate the ISR also increased the formation of subcellular foci of eIF2Bα and eIF2Bε, which differ in their size distribution. Induction of the ISR mainly increased small eIF2Bε punctate bodies, whereas it caused an increase in the formation of eIF2Bα foci of all possible sizes. This change in foci formation by the eIF2B subunits is not due to an underlying change in protein expression (Fig. 2 & S1).

The detection of eIF2Bα foci in the absence of eIF2Bγ suggests the existence of subcellular pools of eIF2Bα dimers which are available to mediate the assembly of eIF2Bβδγε tetramers into full eIF2B(αβδγε)_2_ holocomplexes. Taken together with our previous results showing the link between eIF2B body size and subunit composition and cell type variation in eIF2B bodies (Hodgson et al., 2019; Hanson et al., 2024), our results suggest that a cell–specific mechanism controls the formation of cytoplasmic foci containing various eIF2B subunits which is modulated by the ISR.

### The requirement for eIF2Bα in the formation of eIF2B bodies in mammalian cells

We hypothesised that siRNA inhibition of *EIF2B1* expression would lead to the formation of unassembled eIF2Bβδγε tetramers which cannot assemble a full eIF2B body, thus leading to the reduction in foci larger than 1µm^2^, which was observed (Fig. 3). ISRIB which has been shown to promote the binding of these free tetramers to form functionally active octamers, is able to restore the formation of large eIF2B bodies in neuronal and glial cell lines. We hypothesise that large bodies that form under these conditions are composed of two eIF2Bβγδε tetramers bound by ISRIB, and as such will be unable to respond to stress appropriately

We present compelling evidence pointing towards a crucial necessity of eIF2Bα for the stabilization of the decamer or its subcomplexes which is linked to the formation of eIF2B bodies in mammalian cells. The increase of foci containing isolated eIF2B subunits following cellular stress could provide additional pools of subunits to be incorporated into full eIF2B decamers or alternatively provide catalytic subcomplexes able to perform a lower level of GEF activity to allow translation initiation which is not inhibited by P-eIF2α. This has already been suggested by our identification of foci containing eIF2Bγ and eIF2Bε which have been shown by FRAP analysis to shuttle eIF2α and thereby potentially carry out guanine exchange on eIF2 (Hodgson et al., 2019). Intriguingly, it has recently been proposed that the eIF2Bα homodimer is a sensor of energy modulating the formation of the eIF2B decamer in the presence of specific sugar metabolites (Hao et al 2021). eIF2Bα foci could provide a pool which senses the changes in metabolite levels, thus promoting the formation of the full eIF2B decamer.

Reduction of eIF2Bα expression induces the ISR without the formation of P-eIF2α (Fig. 4). This agrees with studies in mammalian cells showing that the reduction of expression of eIF2Bα triggers the ISR due to the accumulation of unassembled eIF2Bβδγε tetramers (Schoof et al., 2021). .We postulate that endogenous eIF2B(βδγε)_2_ octamers could potentially be altered by the availability of eIF2Bα_2_ homodimers.

### Control of eIF2B subcellular localisation by ISR modulation

We next observed that eIF2B subunits are partially sequestered into SGs upon ISR induction. The proportion of eIF2Bα localisation to SG is higher than that of eIF2Bε. Further analysis of the colocalisation of eIF2B subunits with SG by Airyscan showed that eIF2Bα localises throughout the core and shell of SG whilst eIF2Bε colocalises primarily to the outer shell area of SG (Fig. 5). The functional presence of the eIF2B subunits within SGs is still to be defined fully, however it is of interest to also note that there is preliminary evidence that active translation of mRNA occurs in SG (Mateju et al., 2020). Therefore, the presence of eIF2B catalytic activity may be required to facilitate the initiation of these translating mRNAs.

To elucidate the mechanism causing the relocation of eIF2B subunits to SGs we investigated whether different stress pathways lead to the same level of colocalisation. Inhibition of the eIF4F complex presents an alternative strategy to impede translation initiation and instigate the formation of SGs. Specific chemicals, such as hippuristanol, can obstruct eIF4A’s RNA-binding ability, while others like pateamine A and rocaglamide A (RocA) can induce clamping onto RNA and deplete eIF4A from the eIF4F complex, leading to SG formation through an eIF2α-independent route (Dang et al., 2006; Emara et al., 2010, 2012; Shen & Pelletier, 2020). Hydrogen peroxide (H_2_O_2_) facilitates the binding of 4E-BP1 to eIF4E by promoting a state of hypo-phosphorylation in 4E-BP1, thus reducing translation initiation (Emara et al., 2012) in a P-eIF2α independent manner. Induction of SG by RocA or H_2_O_2_ treatment, without the concomitant formation of P-eIF2α, abolishese colocalisation of eIF2Bα to SG. Our data supports the concept that eIF2Bα plays a pivotal regulatory role of the ISR via the sensing of eIF2α phosphorylation and this direct contact may lead to the incorporation of eIF2Bα into SG.

To further answer this question, we utilised EIF2B1^WT/L99P^ cells which are unable to activate the ISR (Powers, et al., unpublished data). Therefore, we assume that these cells do not have reduced TC levels in response to ER or oxidative stress. This in turn impairs the formation of SGs. Even in the presence of reduced numbers of SG in EIF2B1^WT/L99P^ cells, there is a fall in the proportion of SG which contain colocalised eIF2Bα (Fig. 6).

Our results show that the knockdown of *EIF2B1* expression induces the ISR and formation of SGs without the requirement for phosphorylation of eIF2α. This ISR induction is reversed by treatment with ISRIB in the absence of eIF2Bα expression, most likely due to stabilisation of active eIF2B(βγδε)_2_ structures increasing the amount of ternary complex formation (Figure 4C). Whether eIF2Bα is required for SG formation or SG formation is induced by reduced TC levels cannot be determined from our experimental data. However, it is interesting to note that eIF2Bα knockdown results in inhibition of translation to the same degree as Tg, however the level of SG formation following *EIF2B1* knockdown is lower than that induced by Tg treatment (compare Fig. 7B with Fig. 4C), suggesting that eIF2Bα plays a role in SG formation independent to its role within the eIF2B complex. Recent analysis by Škapik et al., 2025 has shown that the ability of eIF2Bα to sense the activation of the ISR is an essential function enabling colorectal cancer (CRC) cells to continue to proliferate in the presence of increased levels of P-eIF2α. These studies show that reduction of *EIF2B1* expression or inhibition of eIF2Bα dimerisation alters the translation of growth-promoting mRNAs in CRC. We postulate that subcellular localisation of eIF2Bα also contributes to the cellular response to the ISR and the ability of cells to appropriately sense and respond to the presence of increased levels of P-eIF2α. It would be interesting to test this hypothesis by examining SG formation in the presence of gcn-mutations in eIF2Bδ.

In summary, we have identified that eIF2B subunits differ in their association and localisation, potentially affecting the proportion of eIF2B subcomplexes, in a cell type and ISR manner. This is not fully explained by changes in eIF2B subunit expression. We have also shown that individual eIF2B subunits can localise to SGs, which may then impact the ability of cells to form certain subcomplexes of eIF2B. We plan to investigate further whether cell specific binding partners and / or post translational modifications control eIF2B subunit localisation.

## Acknowledgements

This work was funded via a Vice Chancellor Studentship aswwared to Madalena. KO was funded vua a SHRIF funded postdoc.

We thank Dr Paul Clarke & Dr Marissa Powers (ICR) for the SKOV3 parental and SKOV3^WT/L100P^ cell lines.

## Statements and Declarations

The authors have no relevant financial or non-financial interests to disclose

## MATERIALS AND METHODS

### Cell lines and maintenance

### List of reagents and materials

| Reagents | Supplier | Catalog number | Other information |
| --- | --- | --- | --- |
| Minimum Essential Medium (MEM) | Gibco | 11095-080 | 500mL |
| Dulbecco's modified Eagle's medium:F-12 (1:1) (DMEM:F-12) | Gibco | 11574546 |  |
| Dulbecco's Modified Eagle Medium (DMEM) | Gibco | 11574486 | 500mL |
| Rosewell Park Memorial Institute (RPMI) 1640 Medium | Gibco | 11875093 | 500mL |
| Heat-inactivated fetal bovine serum (FBS) | Gibco | 10082-147 | 500mL |

Human glioblastoma astrocytoma cell line (U373-MG) (purchased from ECACCT, #08061901) were cultured in MEM, supplemented with 10% (v/v) FBS, 1% (w/v) NEAA, 1% (v/v) sodium pyruvate, 1% (v/v) L-glutamine and 1% (v/v) P/S. Human glial oligodendrocytic hybrid cell line (MO3.13) (a kind gift from Prof Nicola Woodroofe, originally derived from Cedarlane, #CLU301) were cultured in high glucose DMEM, supplemented with 10 % (v/v) FBS, 1% (v/v) L-glutamine, and 1% (w/v) P/S. Human adrenergic neuroblastoma cell line (SH-SY5Y) (purchased from ATCC, CRL-2266) were cultured in DMEM:F-12 supplemented with 10% (v/v) FBS, 1% (v/v) L-glutamine and 1% (v/v) P/S. All three cell lines were validated with antibodies against lineage-specific markers (data not shown).

Human ovarian adenocarcinoma cell line (SKOV3) and its derived sublines (a kind gift from Prof Paul Clarke, originally purchased from ATCC, HTB-77) were cultured in RPMI, supplemented with 10% (v/v) FBS, 1% (v/v) L-glutamine, and 1% (v/v) P/S. All four cell lines were maintained at 37°C under 5% CO2 and were routinely tested for contamination with MycoAlert^TM^ Mycoplasma Detection Kit purchased from Lonza (LT07-318)

### Cell treatments

For ER stress induction, cells were treated with 1µM Tg for 1h. For oxidative stress induction, cells were treated with 125 μM SA for 30 minutes, 500 μM SA for 1 hour or 0.5 mM, 1 mM or 2 mM H_2_O_2_ for 1 hour. For eIF4A inhibition, cells were treated with 500 nM RocA for 1 hour. For ISIRB treatments, cells were treated with 200 nM ISRIB for 1h. As control, cells were treated with vehicle solution (DMSO) with the highest volume and treatment duration at 37°C depending on the respective drug experimental setup.

### siRNA mediated silencing of *EIF2B1*

U373-MG, MO3.13 and SH-SY5Y were seeded at an appropriate density and cultured for at least 24 hours before transfection. Once the cells reached approximately 70 % confluency, cells were transfected with a SMARTPool of 4 siRNA sequences targeting *EIF2B1* (ON-TARGETplus Human EIF2B1 siRNA; Horizon Discovery L-020180-00-0005). Transfection was carried out using Lipofectamine™ 3000 reagent in accordance with manufacturer’s instructions. Transfection complexes were prepared in FBS-free media at a molar ratio of 1.5:1 (Lipofectamine [µL] : siRNA [µg]) and incubated at RT for 15 minutes. Media was removed and replaced with complete media following 24 hours of transfection. Cells were incubated for an additional 48, 72, 96 or 120h at 37°C under 5 % CO2. As control, cells were transfected with ON-TARGETplus Non-targeting Control siRNAs (Horizon Discovery D-001810-01-05).

### Immunoblotting

5x10^5^ cells were cultured on 6-well plates. Whole-cell protein lysates were prepared in CelLytic M lysis buffer with 1% protease/phosphatase inhibitors and other supplements (17.5 mM β-glycerophosphate, 1 mM PMSF, 10 mM NaF). Lysates were incubated on ice for 10 min and centrifuged (13,000 rpm, 10 min, 4°C) to remove cellular debris. Protein concentrations were determined with Qubit Protein Assay Kit and subjected to SDS-PAGE electrophoresis. For western blots, samples were run on 7.5 or 10% polyacrylamide gel and transferred using Trans-Blot Turbo Mini-nitrocellulose Transfer packs (Bio-Rad) on a Trans-Blot Turbo apparatus. When necessary, membranes were subjected to Revert Total Protein Stain following manufacturer’s instructions. Membranes were blocked in 5% milk or 5% BSA and probed with primary antibodies diluted in 5% milk or 5% BSA, overnight at 4°C. The following antibodies were used: eIF2α (Abcam ab5369; 1:500), phosho-eIF2α[ser51] (Abcam ab32157; 1:500), GAPDH (Cell Signalling #2118; 1:5,000), β-actin (Cell Signalling #3700; 1:2500), eIF2Bα (Proteintech 18010-1-AP; 1:500), eIF2Bβ (Proteintech 11034-1-AP; 1:500), eIF2Bγ (Santa Cruz sc-137248; 1:500), eIF2Bδ (Proteintech 11332-1-AP; 1:50), eIF2Bε(Sigma-Aldrich HPA064370; 1:500). Membranes were then washed 3 times for 5 min/each in TBST, followed by probing with secondary antibodies diluted in 5% milk or 5% BSA in TBST for 1h at RT: goat-anti-rabbit IRDye 680RD (1:10,000) and goat-anti-mouse IRDye 800CW (1:10,000) and washed 3 times for 5 min/each in TBST. Membranes were visualised and quantified on a LiCor Odyssey Scanner with Image Studio Lite software.

### ATF4 ELISA

The Human ATF4 enzyme-linked immunosorbent assay (ELISA) Kit (Proteintech KE00147), was used to determine levels of ATF4, following manufacturer’s instructions.

### Puromycin incorporation assay

91 μM puromycin dihydrochloride was added to media 5 min prior to harvesting and incubated at 37°C. Cells were washed twice with ice-cold PBS supplemented with 355μM cycloheximide, lysed and immunoblotted as described previously. Primary puromycin-specific antibody (1:500) was used to detect puromycinylated proteins. GAPDH was used as loading control.

### Immunocytochemistry

22 × 22 mm squared glass coverslips (Sigma-Aldrich) were rinsed with 100% IMS (Fisher Scientific), added to 6-well plates, and left until IMS fully evaporated. Cells were seeded and transfected as described previously. U373 and SH-SY5Y cells were fixed in ice-cold methanol (Fisher Scientific) at −20°C for 15 min, and MO3.13 cells in 4% (w/v) paraformaldehyde (Alfa Aesar) at RT for 20 min. For methanol fixation, cell membranes were washed with PBS supplemented with 0.5% (v/v) Tween 20 (PBST), 3 times for 3 min and then blocked in PBS supplemented with 1% (w/v) bovine serum albumin (BSA) for 60 min at RT, or overnight at 4°C, under gentle shaker. For paraformaldehyde fixation, cells were washed 3 times with PBST for 3 min, permeabilized with 0.1% (v/v) X-Triton for 5 min at RT and blocked in 1% (w/v) BSA in PBST for 60 min at RT or overnight at 4°C, under gentle shaker. Cell membranes were probed with primary antibodies diluted in 1% (w/v) BSA in PBS, overnight at 4°C under gentle shaker, as following: eIF2Bα (Proteintech 18010-1-AP; 1:25), eIF2Bβ (Proteintech 11034-1-AP; 1:25), eIF2Bγ (Santa Cruz sc-137248; 1:50), eIF2Bδ (Santa Cruz sc-271332; 1:50), eIF2Bε (Sigma-Aldrich HPA069303; 1:25), G3BP-1 (Abcam ab56574 ;1:100). Cells were then washed 3 times with PBST for 5 min, followed by probing with the appropriate host species Alexa Fluor conjugated secondary antibody, diluted in 1% (w/v) BSA in PBS (1:500), for 60 min at RT. Following secondary antibody incubation, cells were washed with PBST, 3 times for 5 min, and mounted with ProLong Gold Antifade Mountant with DAPI (Invitrogen, Thermo Fisher Scientific). Cells were visualised on a Zeiss LSM 800 confocal microscope.

### Confocal imaging

Imaging was performed using a Zeiss LSM 800 confocal microscope combined with Zeiss ZEN 2.3 (blue edition) software for data processing and analysis. Both 40x and 63× plan-apochromat oil objectives were used and a laser with maximum output of 10 mW at 0.2% (488 nm) and 5.0% (561 nm) laser transmissions. Fluorescence crosstalk was minimal and bleed-through was not observed. Image acquisition was performed by orthogonal projection of a z stack of automatically calculated increments for complete single cell imaging.

### Calculating population of cells with localised eIF2B

Percentages of cells with localised eIF2B foci were observed through the Zeiss ZEN 2.3 (blue edition) software. To determine cells with localised versus dispersed signal, a threshold to authenticate eIF2B foci for each imaging conditions/experiment was analysed using the segment region classes method, set up using the automatic triangle threshold (light regions), with 0 % tolerance, 1 % neighborhood and with holes in segmented objects filled for the appropriate secondary antibodies utilised (488 nm and/or 594 nm). Subsequently, a manual set up was carried out in cases of fluctuations of fluorescence between different captured cells. 0 IDs captured per cell was characterised as a dispersed signal and 1 or more IDs captured were characterised as localised eIF2B signal, i.e., cell with localised foci. The counts of cells were performed by DAPI staining using the images of each region of interest. A population of 100 cells were blindly captured and analysed per replicate. The total number of cells with dispersed or localised signal was converted to percentages.

### Determining co-localisation of antibody staining with eIF2B foci

eIF2B foci were analysed as described and assessed on a body-by-body basis of all detected eIF2B foci per cell. eIF2B foci were classed as positive for co-localisation when the two antibody signals overlapped completely. Additionally, profile and 3D surface profiles were used to create profile/surface plots of protein co-localisation. Two size categories were determined, large foci: ≥ 1 μm2 and small foci: < 1 μm2. The counts were carried out in a population of 30 cells per repeat with at least one foci localised.

### Quantification and statistical analysis

All statistical assessments were made in GraphPad Prism 9.2.0 software, with a significance at p < 0.05. All data is presented as means ± standard errors of the mean (s.e.m.). Data was subjected to the Shapiro-Wilk normality test. The following statistical analyses were used dependent on the data: Ordinary one-way ANOVA test followed by a Tukey’s multiple comparisons test (three or more groups of parametric data); Kruskal-Wallis test followed by a Dunn’s multiple comparisons test (three or more groups of non-parametric data); two-way ANOVA test followed by Tukey’s multiple comparisons test (grouped data); Unpaired t-test (two groups of parametric data). Asterisks indicate respective statistical significance as follows: *p < 0.05; **p < 0.01; ***p < 0.001; and ****p < 0.0001.

---

**Figure S1.**
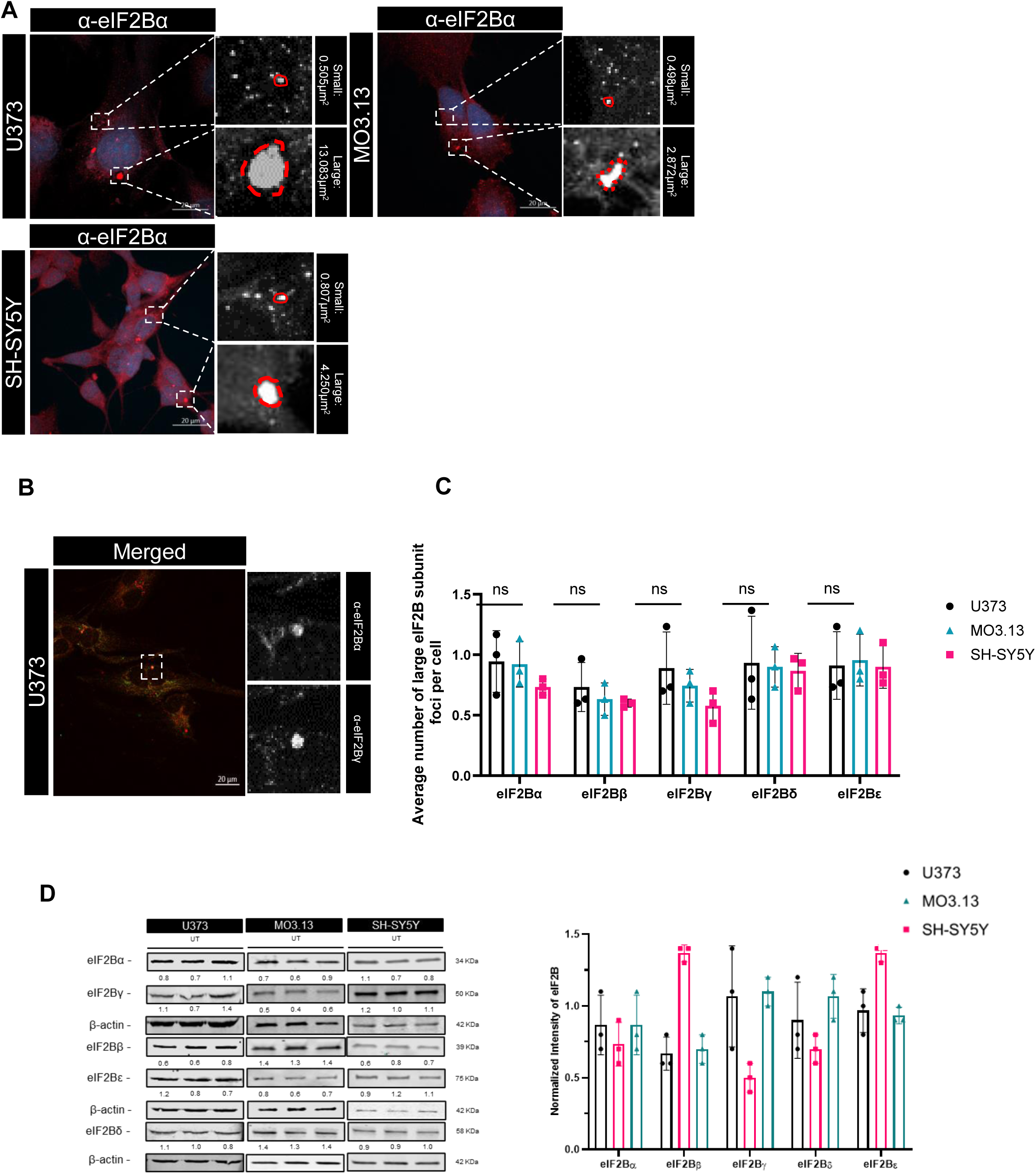
(A) Confocal images of endogenous small and large cytoplasmic foci of eIF2Bα. U373 and SH-SY5Y cells were fixed in methanol, MO3.13 cells were fixed in 4%PFA and subjected to ICC with anti-eIF2Bα primary antibodies and visualized using appropriate secondary antibodies conjugated to Alexa Fluor 594. (B) Confocal images of endogenous eIF2Bα and eIF2Bγ in U373 cells fixed in methanol and subjected to ICC with anti-eIF2Bα and anti-eIF2Bγ primary antibodies visualized using appropriate secondary antibodies conjugated to Alexa Fluor 594 & 488. (C) Average number of large (>1µm^2^) eIF2Bα, eIF2Bβ, eIF2Bγ, eIF2Bδ, or eIF2Bε foci per cell in U373, MO3.13 and SH-SY5Y cells, presented as mean ± SD (n=3 counts of 30 cells with localised foci). *p* Values derived from an ordinary one-way ANOVA test, followed by a Tukey’s multiple analysis, \**p* ≤ 0.05. (D) Western Blot analysis of the levels of eIF2Bα-ε expression in U373, MO3.13 and SH-SY5Y cells under untreated conditions. Levels of eIF2Bα-ε were normalized to levels of β-actin. The average ratio of eIF2Bα protein to β-actin is shown under each cell line, standard deviation in brackets

**Figure S2.**
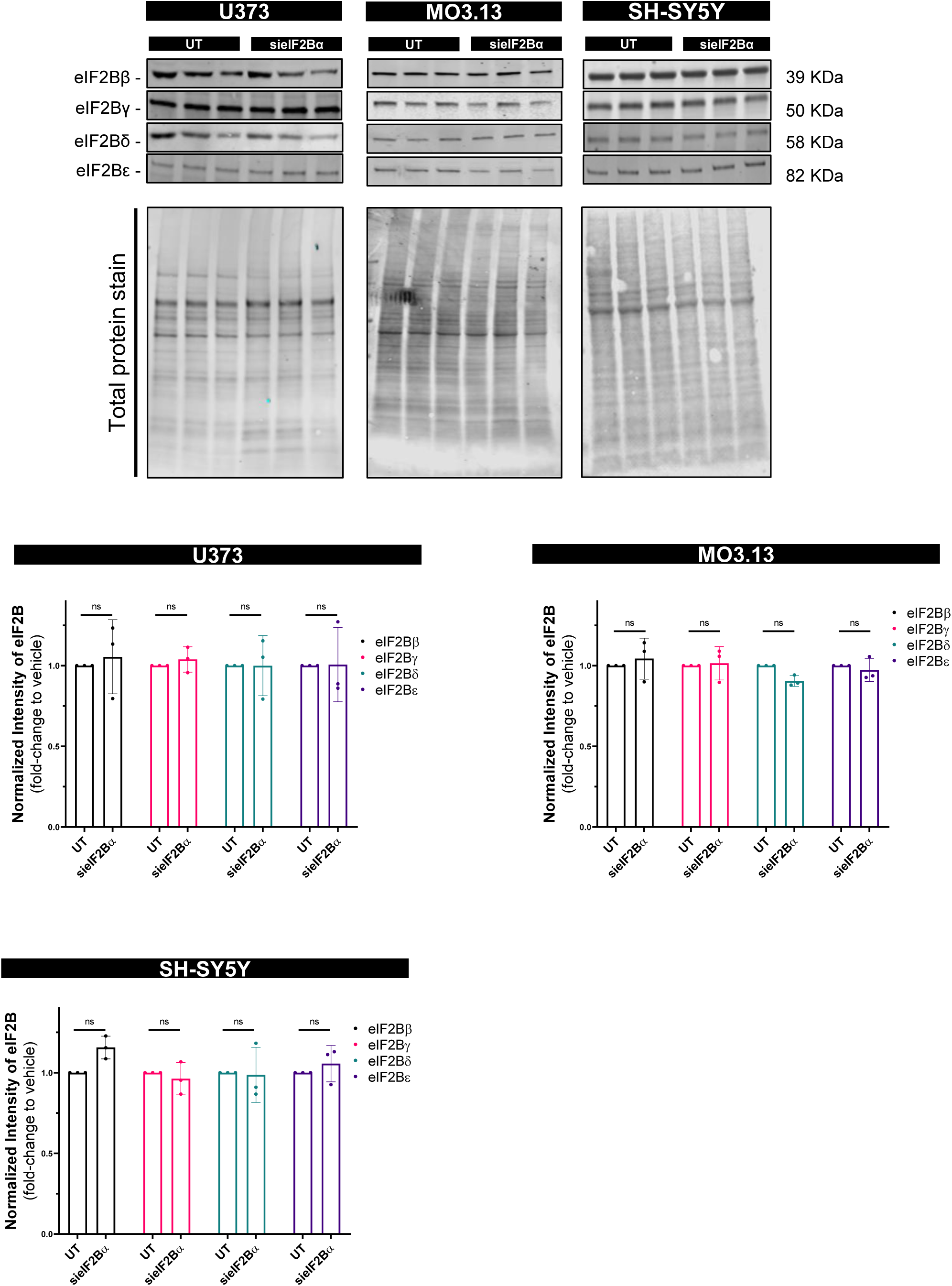
siRNA mediated silencing of EIF2B1 does not alter eIF2Bβ-ε expression levels. Western Blot analysis of the level of eIF2Bββ-ε expression in U373-MG, MO3.13 and SH-SY5Y cells following untreated and siRNA mediated silencing of eIF2Bα for 96h. Levels of eIF2Bββ-ε were normalized to levels of total protein stain and presented as mean ± SD (n=3). Data was analysed by two-way ANOVA test, followed by a Tukey’s multiple analysis. No significant differences observed.

**Figure S3.**
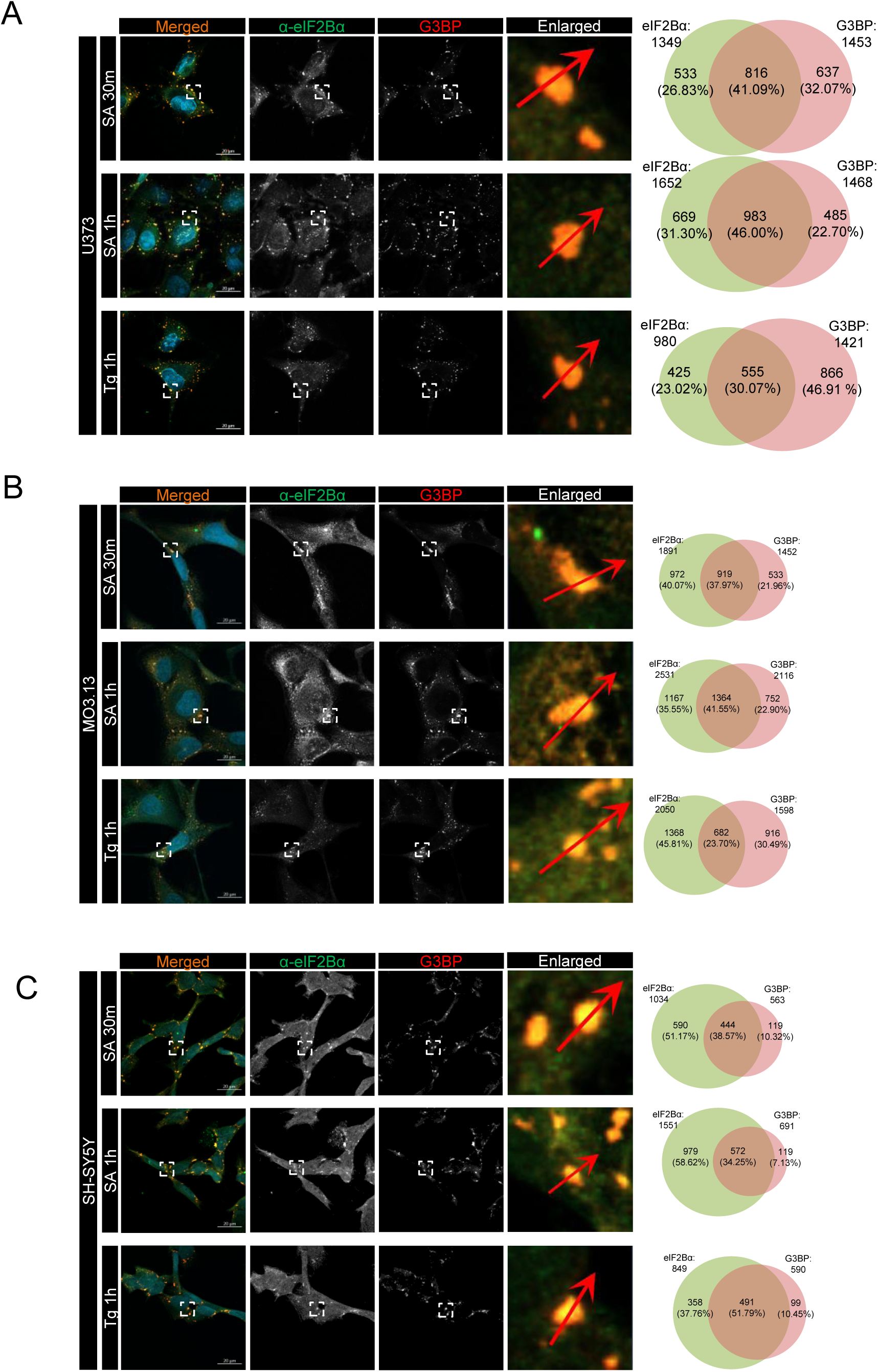
Confocal images of endogenous eIF2Bα and G3BP1 localizing to cytoplasmic foci in cells following treatment with 125 μM SA for 30 minutes, 500 μM SA for 1h or 1 µM Tg for 1h. Cells were subjected to ICC with anti-eIF2Bα (green) and anti-G3BP1 (red primary antibodies and visualized using appropriate secondary antibodies conjugated to Alexa Fluor 488 and 594. (A) U373 cells (B) MO3.13 cells (C) SH SY5Y cells

**Figure S4.**
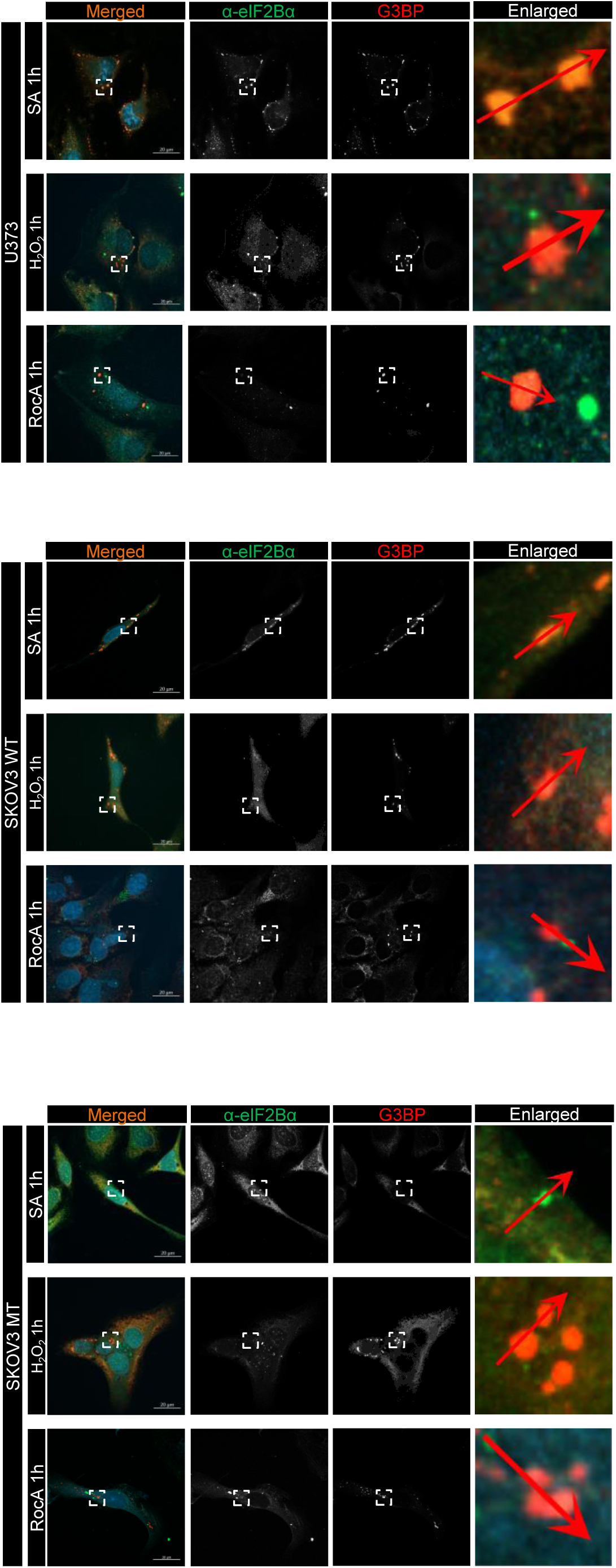
Confocal images of endogenous eIF2Bα and G3BP1 localizing to cytoplasmic foci in U373, SKOV3 EIF2B1 WT/WT and SKOV3 EIF2B1WT/L100P cells following treatment with 500 μM SA for 1h, 0.5mM H_2_O_2_ or 500 nM RocA for 1h. U373 cells were fixed in methanol and subjected to ICC with anti-eIF2Bα (green) and anti-G3BP1 (red primary antibodies and visualized using appropriate secondary antibodies conjugated to Alexa Fluor 488 and 594. The boxed region is enlarged, profile and surface plots were used to show colocalization (separate colours shown on graphics). Percentage of small and large eIF2Bα foci co-localising with G3BP were presented as mean ± SD (n=3 counts in 30 cells with eIF2Bα localisation). Venn diagram of eIF2Bα and G3BP populations and co-localisation (n=3 counts in 30 cells with eIF2Bα localisation).

## Notes

### Competing Interest Statement

The authors have declared no competing interest.

## References

1. Advani, V. M., & Ivanov, P., (2019). Translational control under stress: reshaping the translatome Bioessays 41(5), 1900009

2. Brito Querido, J., Díaz-López, I. & Ramakrishnan, V. The molecular basis of translation initiation and its regulation in eukaryotes. Nat Rev Mol Cell Biol 25, 168–186 (2024). 10.1038/s41580-023-00624-9

3. Campbell, S. G., Hoyle, N. P., & Ashe, M. P. (2005). Dynamic cycling of eIF2 through a large eIF2B-containing cytoplasmic body: implications for translation control. The Journal of Cell Biology, 170(6), 925–934. 10.1083/jcb.200503162

4. Costa-Mattioli, M., & Walter, P. (2020). The integrated stress response: From mechanism to disease. Science (New York, N.Y.), 368(6489), 10.1126/science.aat5314. eaat5314 [pii]

5. De Franco, E., Caswell, R., Johnson, M. B., Wakeling, M. N., Zung, A., Dung, V. C., Bich Ngoc, C. T., Goonetilleke, R., Vivanco Jury, M., El-Khateeb, M., Ellard, S., Flanagan, S. E., Ron, D., & Hattersley, A. T. (2020). De Novo Mutations in EIF2B1 Affecting eIF2 Signaling Cause Neonatal/Early-Onset Diabetes and Transient Hepatic Dysfunction. Diabetes, 69(3), 477–483. 10.2337/db19-1029

6. Dooves, S., Bugiani, M., Postma, N. L., Polder, E., Land, N., Horan, S. T., van Deijk, A. L., van de Kreeke, A., Jacobs, G., Vuong, C., Klooster, J., Kamermans, M., Wortel, J., Loos, M., Wisse, L. E., Scheper, G. C., Abbink, T. E., Heine, V. M., & van der Knaap, M. S. (2016). Astrocytes are central in the pathomechanisms of vanishing white matter. The Journal of Clinical Investigation, 126(4), 1512–1524. 10.1172/JCI83908

7. Egbe, N. E., Paget, C. M., Wang, H., & Ashe, M. P. (2015). Alcohols inhibit translation to regulate morphogenesis in C. albicans. Fungal Genetics and Biology : FG & B, 77, 50–60. 10.1016/j.fgb.2015.03.008

8. Glauninger, H., Wong Hickernell, C. J., Bard, J. A. M., & Drummond, D. A. (2022). Stressful steps: Progress and challenges in understanding stress-induced mRNA condensation and accumulation in stress granules. Molecular Cell, 82(14), 2544–2556. 10.1016/j.molcel.2022.05.014

9. Hanson, F. M., Hodgson, R. E., de Oliveira, M. I. R., Allen, K. E., & Campbell, S. G. (2022). Regulation and function of elF2B in neurological and metabolic disorders. Bioscience Reports, 42(6), BSR20211699. doi: 10.1042/BSR20211699. 10.1042/BSR20211699

10. Hanson, F. M., Ribeiro de Oliveira, M. I., Cross, A. K., Allen, K. E., & Campbell, S. G. (2024). eIF2B localization and its regulation during the integrated stress response is cell-type specific. iScience, 27(9)10.1016/j.isci.2024.110851

11. Hinnebusch, A. G. Y. The Scanning Mechanism of Eukaryotic Translation Initiation. Annual Review of Biochemistry, 83(Volume 83, 2014), 779–812. 10.1146/annurev-biochem-060713-035802

12. Hinnebusch, A. G., Ivanov, I. P., & Sonenberg, N. (2016). Translational control by 5’-untranslated regions of eukaryotic mRNAs. Science (New York, N.Y.), 352(6292), 1413–1416. 10.1126/science.aad9868

13. Hodgson, R. E., Varanda, B. A., Ashe, M. P., Allen, K. E., & Campbell, S. G. (2019). Cellular eIF2B subunit localization: implications for the integrated stress response and its control by small molecule drugs. Molecular Biology of the Cell, 30(8), 942–958. 10.1091/mbc.E18-08-0538

14. Kashiwagi, K., Yokoyama, T., Nishimoto, M., Takahashi, M., Sakamoto, A., Yonemochi, M., Shirouzu, M., & Ito, T. (2019). Structural basis for eIF2B inhibition in integrated stress response. Science (New York, N.Y.), 364(6439), 495–499. 10.1126/science.aaw4104

15. Kedersha, N., Chen, S., Gilks, N., Li, W., Miller, I. J., Stahl, J., & Anderson, P. (2002). Evidence that ternary complex (eIF2-GTP-tRNA(i)(Met))-deficient preinitiation complexes are core constituents of mammalian stress granules. Molecular Biology of the Cell, 13(1), 195–210. 10.1091/mbc.01-05-0221

16. Kenner, L. R., Anand, A. A., Nguyen, H. C., Myasnikov, A. G., Klose, C. J., McGeever, L. A., Tsai, J. C., Miller-Vedam, L. E., Walter, P., & Frost, A. (2019). eIF2B-catalyzed nucleotide exchange and phosphoregulation by the integrated stress response. Science (New York, N.Y.), 364(6439), 491–495. 10.1126/science.aaw2922

17. Kuhle, B., Eulig, N. K., & Ficner, R. (2015). Architecture of the eIF2B regulatory subcomplex and its implications for the regulation of guanine nucleotide exchange on eIF2. Nucleic Acids Research, 43(20), 9994–10014. 10.1093/nar/gkv930

18. Leferink, P. S., Dooves, S., Hillen, A. E. J., Watanabe, K., Jacobs, G., Gasparotto, L., Cornelissen-Steijger, P., van der Knaap, M. S., & Heine, V. M. (2019). Astrocyte Subtype Vulnerability in Stem Cell Models of Vanishing White Matter. Annals of Neurology, 86(5), 780–792. 10.1002/ana.25585

19. Marintchev, A., & Ito, T. (2020). eIF2B and the Integrated Stress Response: A Structural and Mechanistic View. Biochemistry, 59(13), 1299–1308. 10.1021/acs.biochem.0c00132

20. Mateju, D., Eichenberger, B., Voigt, F., Eglinger, J., Roth, G., & Chao, J. A. (2020). Single-Molecule Imaging Reveals Translation of mRNAs Localized to Stress Granules. Cell, 183(7), 1801–1812.e13. 10.1016/j.cell.2020.11.010

21. Norris, K., Hodgson, R. E., Dornelles, T., Allen, K. E., Abell, B. M., Ashe, M. P., & Campbell, S. G. (2020). Mutational analysis of the alpha subunit of eIF2B provides insights into the role of eIF2B bodies in translational control and VWM disease. The Journal of Biological Chemistry, jbc.RA120.014956 [pii]

22. Nuske, E., Marini, G., Richter, D., Leng, W., Bogdanova, A., Franzmann, T. M., Pigino, G., & Alberti, S. (2020). Filament formation by the translation factor eIF2B regulates protein synthesis in starved cells. Biology Open, 9(7), 10.1242/bio.046391. bio046391 [pii]

23. Reid, P. J., Mohammad-Qureshi, S., & Pavitt, G. D. (2012). Identification of Intersubunit Domain Interactions within Eukaryotic Initiation Factor (eIF) 2B, the Nucleotide Exchange Factor for Translation Initiation*. Journal of Biological Chemistry, 287(11), 8275–8285. 10.1074/jbc.M111.331645

24. Schoof, M., Boone, M., Wang, L., Lawrence, R., Frost, A., & Walter, P. (2021). eIF2B conformation and assembly state regulates the integrated stress response. eLife, 10, 10.7554/eLife.65703. 10.7554/eLife.65703

25. Škapik, I. P., Giacomelli, C., Hahn, S., Deinlein, H., Gallant, P., Diebold, M., Biayna, J., Hendricks, A., Olimski, L., Otto, C., Kastner, C., Wolf, E., Schülein-Völk, C., Maurus, K., Rosenwald, A., Schleussner, N., Jackstadt, R., Schlegel, N., Germer, C., . . . Wiegering, A. (2025). Maintenance of p-eIF2α levels by the eIF2B complex is vital for colorectal cancer. The EMBO Journal,, 1–31; 31. 10.1038/s44318-025-00381-9

26. Triñanes-Ramos, J., Bugiani, M., van Rooijen-van Leeuwen, G., Chevalier, J. A., Siu, Y. J., van Utenhove, E.L.H., Hoogterp, L., Witkamp, D., Oudejans, E., Lodder, B., Kole, M., Saher, G., Nave, K., Abbink, T. E. M., & van der Knaap, M.S. (2025). Cell-specific Eif2b5 mutant mice: novel insights into roles of macroglia in vanishing white matter. Brain, awaf171. 10.1093/brain/awaf171

27. Wortham, N. C., & Proud, C. G. (2015). eIF2B: recent structural and functional insights into a key regulator of translation. Biochemical Society Transactions, 43(6), 1234–1240. 10.1042/BST20150164

28. Zyryanova, A. F., Kashiwagi, K., Rato, C., Harding, H. P., Crespillo-Casado, A., Perera, L. A., Sakamoto, A., Nishimoto, M., Yonemochi, M., Shirouzu, M., Ito, T., & Ron, D. (2021). ISRIB Blunts the Integrated Stress Response by Allosterically Antagonising the Inhibitory Effect of Phosphorylated eIF2 on eIF2B. Molecular Cell, 81(1), 88–103.e6. S1097-2765(20)30738-3 [pii]

